# “Monoclonal-type” plastic antibodies for SARS-CoV-2 based on Molecularly Imprinted Polymers

**DOI:** 10.1101/2020.05.28.120709

**Authors:** Ortensia Ilaria Parisi, Marco Dattilo, Francesco Patitucci, Rocco Malivindi, Vincenzo Pezzi, Ida Perrotta, Mariarosa Ruffo, Fabio Amone, Francesco Puoci

## Abstract

Our idea is focused on the development of “monoclonal-type” plastic antibodies based on Molecularly Imprinted Polymers (MIPs) able to selectively bind a portion of the novel coronavirus SARS-CoV-2 spike protein to block its function and, thus, the infection process. Molecular Imprinting, indeed, represents a very promising and attractive technology for the synthesis of MIPs characterized by specific recognition abilities for a target molecule. Given these characteristics, MIPs can be considered tailor-made synthetic antibodies obtained by a templating process.

In the present study, the developed imprinted polymeric nanoparticles were characterized in terms of particles size and distribution by Dynamic Light Scattering (DLS) and the imprinting effect and selectivity were investigated by performing binding experiments using the receptor-binding domain (RBD) of the novel coronavirus and the RBD of SARS-CoV spike protein, respectively. Finally, the hemocompatibility of the prepared MIP-based plastic antibodies was also evaluated.

## Main Text

Coronaviruses (CoVs) are single-stranded enveloped positive-sense RNA viruses belonging to the subfamily *Coronavirinae*, family *Coronavirdiae*, and characterized by a RNA genome ranging from 26 to 32 kilobases (Chan et al., 2020; Su et al., 2016). On the basis of their host specificity, these CoVs can be classified into four main genera such as *Alphacoronavirus* (αCoV), *Betacoronavirus* (βCoV), *Deltacoronavirus* (δCoV) and *Gammacoronavirus* (γCoV) (Snijder et al., 2006). Coronaviruses have been identified in birds and mammals such as bats, mice, cats and dogs (Fehr and Perlman, 2015; Lu et al., 2020), but this kind of viruses is able to infect both animals and humans. Human CoVs, indeed, were first characterized in the 1960s and are associated with respiratory infections (Paraskevis et al., 2020). In December 2019, several cases of patients affected by viral pneumonia were reported in Wuhan City, Hubei Province, China and a novel coronavirus (SARS-CoV-2, initially called 2019-nCoV) able to infect humans was identified and detected in these patients.

The genome analysis of the novel coronavirus revealed that it belongs to the subgenus *Sarbecovirus* of the genus *Betacoronavirus* and it is closely related (88% identity) to two bat-SARS-like coronavirus (bat-SL-CoVZC45 and bat-SL-CoVZXC21) with which it forms a distinct lineage (Lu et al., 2020; Pradhan et al., 2020). On the other hand, SARS-CoV-2 is divergent from SARS-CoV (about 79% similarity) and MERS-CoV (about 50% similarity) (Lu et al., 2020). The genome of SARS-CoV-2 encodes different proteins such as the spike protein, the envelope protein, the membrane protein, and the nucleocapsid protein. The coronavirus spike protein is a surface protein that mediates host recognition and attachment. It consists of two functional subunits: the S1 subunit, which contains a receptor-binding domain (RBD) responsible for host cell receptor recognizing and binding, and the S2 subunit, which is involved in the viral and host membranes fusion. These two processes, which represent the initial steps in the coronavirus infection cycle, are crucial in determining host specificity, tissue tropism and transmission capacity (Lu et al., 2015; Wang et al., 2016). Although the SARS-CoV-2 genome was closer to bat-SL-CoVZC45 and bat-SL-CoVZXC21, its RBD structure presents a high homology to that of SARS-CoV, which uses angiotensin-converting enzyme 2 (ACE2) as host cell receptor (Li et al., 2005). Both SARS-CoV and SARS-CoV-2 belong to the β-genus and, in a recent study (Wan et al., 2020), it is reported that the overall sequence similarities between 2019-nCoV and SARS-CoV spike proteins are around 76%-78% for the whole protein and around 73%-76% for the RBD. Moreover, Xu X. et al. (Xu et al., 2020) have found that the novel coronavirus spike protein has a relevant binding affinity to human ACE2 and, thus, this virus can interact with this entry receptor causing the infection of human respiratory epithelial cells. Therefore, the spike protein, which is involved in viral recognition and binding to human ACE2 (Lei et al., 2020; Yan et al., 2020; Zhou et al., 2020) playing a key role also in human-to-human transmission of this novel coronavirus, represents the common and primary target for the development of antibodies, vaccines and therapeutic agents.

In this context, our idea is to develop “monoclonal-type” plastic antibodies based on Molecularly Imprinted Polymers (MIPs) for the selective recognition and binding of the RBD of the novel coronavirus SARS-CoV-2 in the aim to block the function of its spike protein (*Figure 1*.).

**Figure 1.**
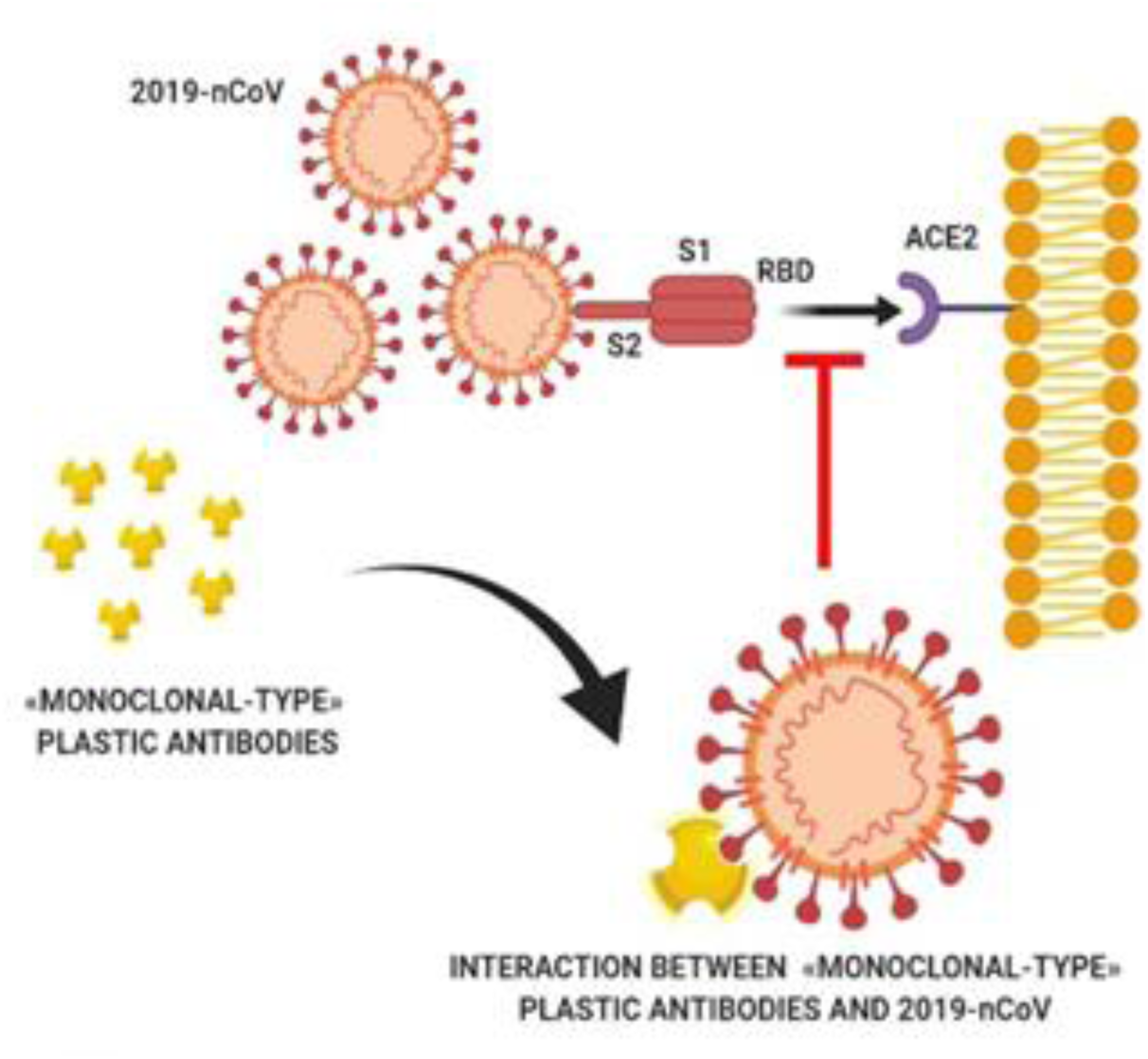
Schematic representation of the interaction between MIP-based “monoclonal-type” plastic antibodies and SARS-CoV-2.

This kind of polymeric matrices is synthesized by polymerizing functional and crosslinking monomers around the template (Parisi et al., 2014; Parisi et al., 2020a). The selective recognition abilities of the imprinted polymers are due to the formation of a complex between the target analyte and the selected functional monomers during the pre-polymerization step. Therefore, the choice of the monomers represents a crucial point in the preparation of effective MIPs and it is based on their ability to establish interactions with the functional groups of the template molecule in a covalent or non-covalent way. Three main approaches, indeed, can be used to synthesize this kind of polymers depending on the nature of the interactions occurring between the template molecule and the chosen functional monomers during both pre-polymerization and binding steps (Parisi et al., 2020b; Puoci et al., 2010). In the covalent one, template and functional monomers are covalently bound during the pre-polymerization phase and, after the polymerization reaction, the analyte is extracted from the polymeric matrix by chemical cleavage of the covalent bonds. Then, the same covalent interactions are re-formed during the rebinding. The semi-covalent approach involves the formation of covalent interactions during the polymerization process and non-covalent interactions in the rebinding process. Finally, the non-covalent approach is based on the formation of non-covalent interactions, including hydrogen bonds and electrostatic, π-π and hydrophobic interactions, between template and monomers during the polymerization process and the subsequent recognition phase. This method is widely employed due to several advantages such as the simple experimental procedure and the large variety of appropriate functional monomers. In the present study, the last imprinting approach was chosen for the MIPs-based antibodies preparation due to the nature of the template-monomers interactions, which are similar to those found in biological systems. Once the polymerization reaction has taken place, the template is extracted leading to a porous crosslinked polymeric matrix containing binding holes fitting size, shape and functionalities of the target compound.

Plastic antibodies made from tailor-made polymeric imprinted nanoparticles represent an alternative to the traditional antibodies, which require an expensive production procedure and are often unreliable due to their restricted stability (Refaat et al., 2019; Xu et al., 2019). Being synthetic materials, MIPs are robust, physically and chemically stable in a wide range of conditions, including temperature and pH, and more easily available due to a low-cost, reproducible and relatively fast and easy preparation compared to the biological counterpart (Piloto et al., 2018; Wubulikasimu et al., 2019). Therefore, this kind of materials combines the robustness of polymers with the selectivity of natural receptors (Capriotti et al., 2020) and could find applications in several fields, including separation, catalysis and drug delivery, as sensors, synthetic biomimetic receptors and recognition elements in bioanalytical assays (Pan et al., 2018). Moreover, imprinted polymers are characterized by significant versatility. These polymeric materials, indeed, can be designed and engineered according to their specific application developing polymers with magnetic and/or fluorescent properties.

Synthetic polymeric antibodies against SARS-CoV-2 was produced according to the Molecular Imprinting Technology (MIT) and a Non-Imprinted Polymer (NIP) was also prepared following the same experimental procedure adopted for the imprinted nanoparticles, but in the absence of the novel coronavirus RBD. For this purpose, the non-covalent imprinting approach was adopted and biocompatible functional and crosslinking monomers were chosen.

Particles size and distribution were investigated by Dynamic Light Scattering (DLS) using a Zetasizer nano ZS DLS (Malvern Instrument, Malvern, UK). For sample preparation, 25 μL of a polymeric nanoparticles suspension in phosphate buffer solution (PBS, 10^−3^ M) at pH 7.4 were added to 3 mL of PBS and sonicated for 30 seconds. The performed DLS analysis highlighted a mean diameter equal to 80.3 ± 6.2 nm and a polydispersity index (PI) of 0.287 indicating a narrow size distribution (*Figure 2*.). The dimensionless polydispersity index, indeed, was used as a measure of the size distribution and a value below 0.3 is indicative of a homogeneous population of particles.

**Figure 2.**
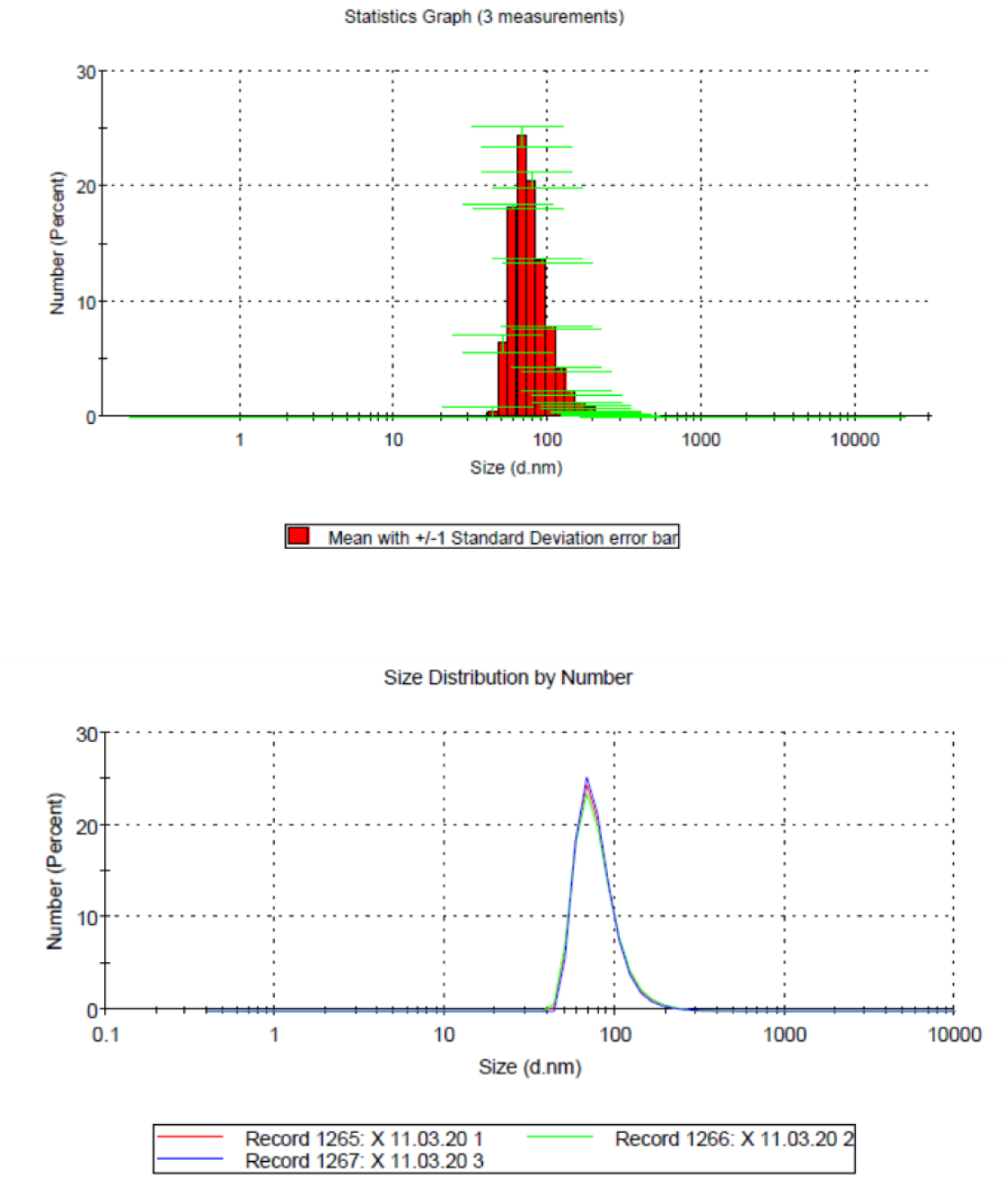
Characterization of Molecularly Imprinted Nanoparticles: DLS analysis.

In order to investigate both recognition properties and selectivity of the prepared imprinted nanoparticles, binding studies were carried out in phosphate buffer solution (PBS) at pH 7.4. The imprinting effect was evaluated by binding experiments in which amounts of imprinted and non-imprinted nanoparticles were incubated with a standard solution of the SARS-CoV-2 receptor-binding domain. SDS-PAGE Electrophoresis was used to quantify the interaction of MIP with the novel coronavirus RBD and the obtained results (*Figure 3*. and *Table 1*.) highlighted the capability of the imprinted polymer to recognize and bind a higher amount of the target molecule compared to the corresponding non-imprinted one.

**Table 1.** Evaluation of the Imprinting Effect and Selectivity. Percentages of bound SARS-CoV-2 receptor-binding domain and SARS-CoV receptor-binding domain by imprinted (MIP) and non-imprinted (NIP) nanoparticles. Data are shown as means ± S.D.

| BOUND SARS-CoV-2 RBD (%) |  | BOUND SARS-CoV RBD (%) |  |
| --- | --- | --- | --- |
| MIP | NIP | MIP | NIP |
| 29.30 ± 1.13 | 3.70 ± 0.58 | 0.68 ± 0.03 | 0.17 ± 0.08 |

**Figure 3.**
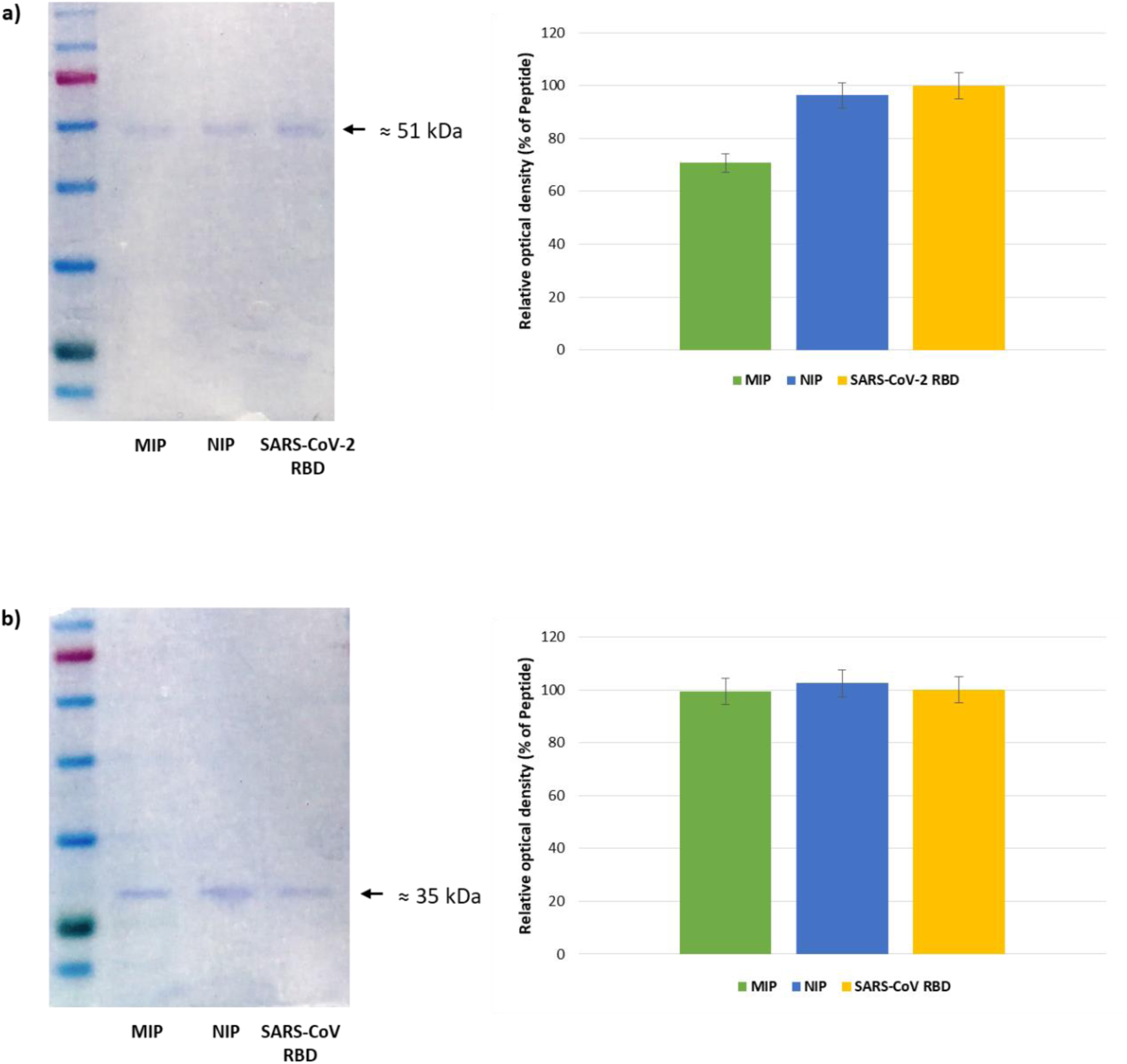
Binding Studies - SDS-PAGE Electrophoresis (Coomassie blue stained): **a)** SARS-CoV-2; **b)** SARS-CoV.

The same experimental conditions were adopted to perform selectivity studies, which involved a standard solution of a molecule structurally similar to the 2019-nCoV RBD such as the RBD of SARS-CoV spike protein. The obtained results showed no significant differences between MIP and NIP nanoparticles (*Table 1*.) in the interaction with SARS-CoV RBD confirming the specific and selective abilities of the imprinted polymer. These MIP properties, indeed, are due to the presence of selective molecular recognition holes that are complementary to the target template in terms of size, shape and functional groups. The choice of functional monomer plays a key role in the imprinting process to ensure the formation of selective binding sites within the polymeric matrix. In this work, the employed functional monomer was chosen in the aim to promote also hydrophobic interactions with the receptor-binding domain of the novel coronavirus. Therefore, the interactions established between the SARS-CoV-2 RBD and the synthesized imprinted polymeric nanoparticles are not affected by pH changes.

The developed MIP-based plastic antibodies were designed for intravenous administration and, therefore, have to provide a suitable hemocompatibility, which is a key requirement for a successful system that comes in contact with the bloodstream. In order to evaluate the hemocompatibility of the prepared imprinted nanoparticles, hemolysis tests were performed using an isotonic phosphate buffer solution and pure water as negative and positive controls, respectively. A polymeric material is considered not-hemolytic if the hemolysis percentage is below 5% (Contreras-García et al., 2011; Panikkar et al., 1997). The performed hemolysis assay revealed that the synthesized imprinted nanoparticles induced 3.9% hemolysis, thus, the observed hemolytic potential is within satisfactory limits indicating a good hemocompatibility without causing erythrocyte damage.

In conclusion, Molecular Imprinting Technology was adopted as synthetic strategy to prepared molecularly imprinted nanoparticles able to selectively recognize and bind the spike protein receptor-binding domain of the novel coronavirus SARS-CoV-2. The reported preliminary results suggested the potential use of this biocompatible polymeric material as MIP-based “monoclonal-type” plastic antibodies devoted to block the function of the virus spike protein. Given these characteristics, the developed nanoparticles could be potentially used as free-drug therapeutics in the treatment of 2019-nCoV infection. Moreover, when loaded with antiviral agents, these nanoparticles could act as a powerful multimodal system combining their ability to block the virus spike protein with the targeted delivery of the loaded drug. In addition, the same nanoparticles can be further engineered to become an immunoprotective vaccine or a MIP-based sensor for diagnostic purpose.

## Acknowledgments

This work was supported by University of Calabria and Macrofarm s.r.l., a spin-off company of the University of Calabria.

## Author contributions

Direction of the Project, F.P.*; Conceptualization, Supervision and Data Curation, F.P.*, O.I.P. and V.P.; Methodology and Formal Analysis, M.D., R.M., F.P., M.R., I.P. and F.A.; Writing, O.I.P.

